# Trehalose 6-phosphate activates Target of Rapamycin in plants

**DOI:** 10.64898/2025.12.11.693803

**Authors:** Hui Liu, Jantana Blanford, Hai Shi, Saroj Sah, Sanket Anaokar, Jorg Schwender, John Shanklin, Zhiyang Zhai

## Abstract

Coordinating carbon availability with growth is a fundamental challenge in plants. Target of Rapamycin (TOR) promotes cell growth and is activated by sugars, but the underlying molecular mechanism has remained elusive. Here, we identify trehalose 6-phosphate (T6P), a sucrose-derived metabolite, as the key signal linking carbon to TOR activity. In Arabidopsis and *Brassica napus*, T6P stimulates cell growth via TOR and is required for sucrose-induced TOR activation. We show immunoprecipitates using anti-TOR antibodies contain catalytically active TOR in addition to the catalytically active SnRK1 energy sensor kinase. *In vitro experiments show that*, SnRK1α1, the catalytic subunit of SnRK1 suppresses TOR activity, and T6P reverses this inhibition in a dose-dependent manner, providing biochemical evidence that T6P activates TOR by suppressing SnRK1. This work thus establishes a direct sucrose–T6P–SnRK1–TOR signaling axis that couples carbon availability to plant growth.

## Introduction

Balancing carbon availability with growth is a central challenge for all organisms. In plants, the kinase Target of Rapamycin (TOR) serves as a master regulator that integrates nutrient and energy signals to promote biosynthesis, cell proliferation, and developmental progression. TOR functions in a conserved complex with RAPTOR (regulatory associated protein of TOR) and LST8 (lethal with SEC13 protein 8), and activates growth by phosphorylating downstream effectors such as the 40S ribosomal protein S6 kinase (S6K), which phosphorylates ribosomal protein S6 (RPS6), thereby stimulating translation and metabolism^1, 2, 3, 4, 5, 6^.

Plant TOR activity is known to be triggered by light, auxin, amino acids, and sugars^1, 7, 8, 9, 10^. While light and hormonal pathways converge on TOR through the Rho-like GTPase ROP2^7, 8, 9^, the mechanism by which sugars activate TOR has remained unresolved. Both glucose and sucrose stimulate TOR, but since these metabolites are rapidly consumed, it has been unclear whether they act as direct signals or through metabolic intermediates. This missing link has left a fundamental gap in understanding how carbon status is sensed and relayed to TOR.

Trehalose 6-phosphate (T6P) is a sucrose-derived metabolite that reflects sucrose availability and regulates plant growth and development^11^. T6P is synthesized by Trehalose 6-phosphate synthase 1 (TPS1) from glucose 6-phosphate and UDP-glucose^12^. In contrast, Sucrose non-fermenting-1-related protein kinase 1 (SnRK1) acts as a central energy sensor, maintaining homeostasis under stress and generally antagonizing growth^13, 14, 15^. SnRK1 and TOR signaling are tightly interconnected and often function antagonistically at the systems level^16, 17^. Under energy deprivation, SnRK1 inhibits TOR, likely through direct phosphorylation of RAPTOR^18, 19^. TOR also interacts with SnRK1 signaling through the SnRK2/PYR/PP2A/ABA pathway^20, 21, 22^. Importantly, T6P inhibits SnRK1 activity in growing tissues via a heat-labile factor^23, 24^, and recent studies indicate that T6P directly binds SnRK1α1, preventing its phosphorylation by Geminivirus Rep Interacting Kinase 1 (GRIK1) and suppressing SnRK1 activation^25, 26, 27, 28^. Consistent with this antagonism, mutations in SnRK1α1 and its regulatory subunit SNF4 rescue embryo lethality and flowering defects of a *tps1* knockout^29^. It was reported that T6P increases root branching through inhibition of SnRK1 and activation of TOR, though coordinated inhibition of SnRK1 and activation of TOR, though whether these changes are related or occur in parallel was not determined^30^. However, the same work showed that activation of TOR by T6P was accompanied by an increase the level of TOR expression and its phosphorylation status. Recently we reported that direct binding of T6P to SnRK1α1 prevents its activation by its upstream activating kinase GRIK1^27^. Here we present evidence that the immunoprecipitated TOR complex contains both active TOR and SnRK1, and that SnRK1 suppresses TOR activity, and T6P reverses this inhibition in a dose-dependent manner, providing biochemical evidence to support a sucrose–T6P–SnRK1–TOR signaling axis connecting carbon availability to TOR activity that drives plant growth.

## Results and Discussion

### T6P and TOR converge to regulate growth pathway

To test whether T6P and TOR act within a shared growth pathway, we examined plant thermomorphogenesis, an adaptive growth response characterized by hypocotyl and petiole elongation at elevated temperature that requires carbon availability^31^. As previously reported, dexamethasone (DEX)-inducible *TPS1* in a *tps1* knockout background (*tps1-2*; *GVG*::*TPS1*; hereafter *tps1*) displayed reduced thermoresponsive growth in the absence of DEX^26^. We confirmed this phenotype; *tps1* seedlings exhibited only a 1.4-fold increase in hypocotyl length at 28 °C compared to 20 °C, whereas wild type (WT) seedlings elongated 5.3-fold under the same conditions (Fig. 1a, Supplementary Fig. 1). To further evaluate the role of T6P in thermomorphogenesis, we analyzed Arabidopsis lines overexpressing *otsA* (encoding *E. coli* T6P synthase) or *otsB* (encoding T6P phosphatase). Overexpression of *otsA* significantly enhanced the thermoresponsive growth, while *otsB* overexpression markedly reduced growth relative to WT (Fig. 1b, Supplementary Fig. 2).We next assessed TOR’s contribution using the estradiol-inducible TOR silencing line (*tor-es1*)^4^. In the absence of estradiol, *tor-es1* resembled WT; however, estradiol treatment reduced hypocotyl elongation at 28 °C by 80% (Fig. 1c). Similarly, treatment of WT seedlings with Torin2, a selective ATP-competitive TOR inhibitor^32^, caused a 32% reduction in thermoresponsive hypocotyl growth compared to untreated control (Supplementary Fig. 3). Microscopy revealed that epidermal cells of hypocotyl were markedly shorter in estradiol-treated *tor-es1* (64% reduction) and Torin2-treated WT (65% reduction) relative to their respective controls (Supplementary Fig. 4). By contrast, two hypermorphic TOR alleles (G548 and S7817) ^33^and *S6K1* overexpression lines exhibited longer hypocotyls at 28 °C (Supplementary Fig. 5). Together, results from these three independent approaches demonstrate that T6P and TOR are essential positive regulators of thermomorphogenesis.

**Fig. 1.**
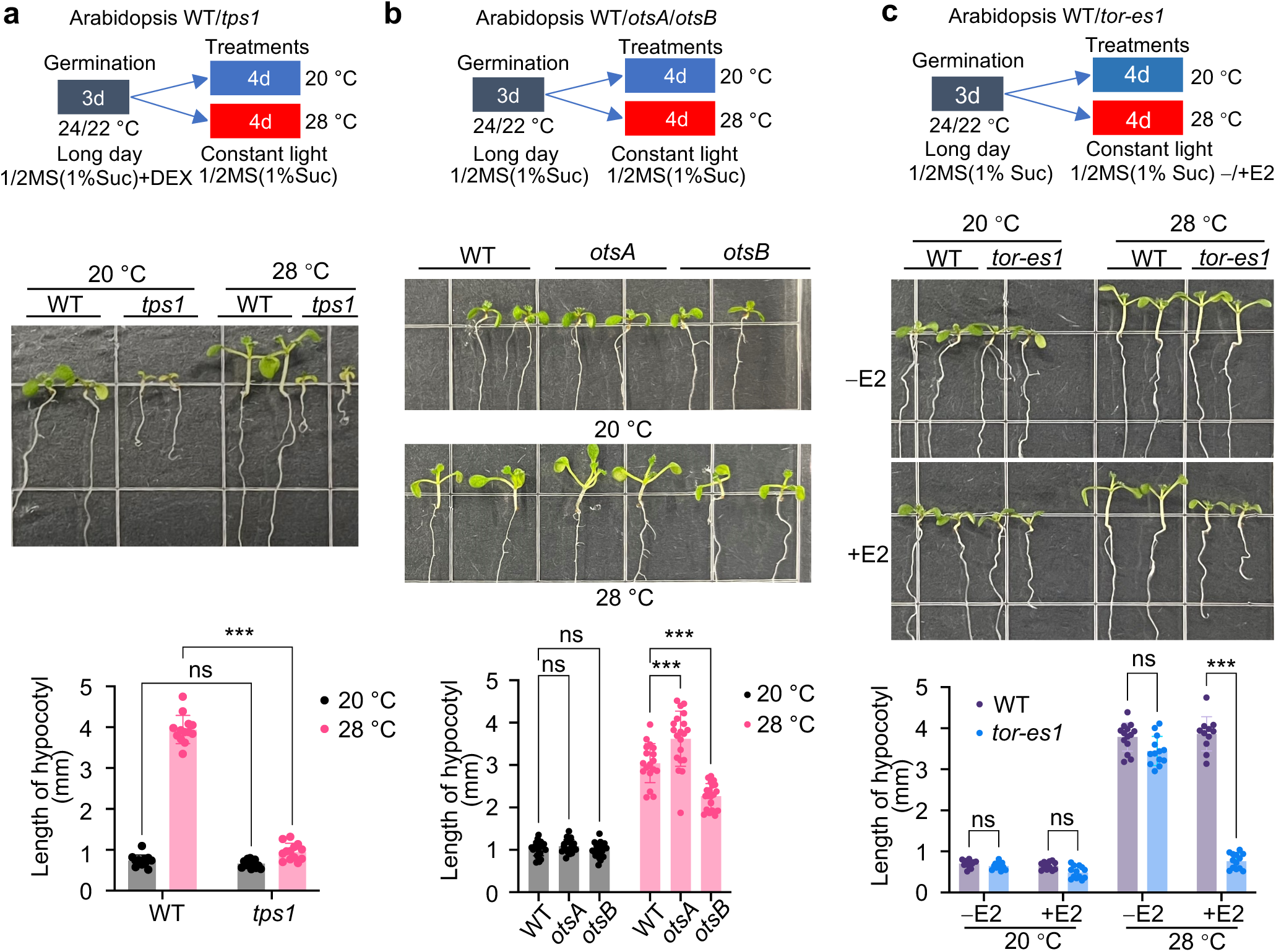
TOR and T6P signaling regulate thermomorphogenesis in Arabidopsis. **a**, Loss of T6P reduces thermoresponsive growth. Hypocotyl elongation is attenuated in *tps1-2*;*GVG*::*TPS1* (hereafter *tps1*) seedlings compared with wild type (WT) at 28 °C. WT and *tps1* were germinated on ½ MS [1% sucrose (Suc)] containing 10 μM dexamethasone (DEX) for 3 d (long-day conditions), then transferred to fresh ½ MS (1% sucrose) and grown at 20 °C or 28 °C for 4 d. Representative seedlings of WT and *tps1* are shown, with quantified hypocotyl lengths below. Grid size, 13.5×13.5mm. In the absence of DEX, *tps1* T6P levels are reduced (see Supplementary Fig. 1). **b**, Altered T6P levels modulate growth responses. *otsA* overexpression increases T6P and enhances hypocotyl elongation, whereas *otsB* overexpression reduces T6P and attenuates growth. T6P measurements are provided in Supplementary Fig. 2. **c**, TOR inhibition diminishes thermoresponses. Estradiol-induced TOR silencing (*tor-es1*) or pharmacological inhibition with Torin2 reduces hypocotyl elongation at 28 °C. WT and *tor-es1* were germinated on ½ MS (1% sucrose) for 3 d (long-day conditions), then transferred to ½ MS (1% sucrose) without (−) or with (+) 1 μM of estradiol (E2) and grown at 20 or 28 °C for 4 d under constant light (30 μmol m⁻² s⁻¹). In all panels, significance by Tukey’s multiple comparisons: *, *P* < 0.05; **, *P* < 0.01; ***, *P* < 0.001, ns, not significant. Together, these data show that T6P and TOR are both required for thermomorphogenesis.

### T6P promotes growth of *Brassica napus* suspension cells in a TOR-dependent manner

To determine whether T6P directly stimulates cellular growth via TOR, we employed *Brassica napus* (*B. napus*) suspension cultures, a system previously shown to import exogenously supplied T6P^25^. Treatment with 0.25 μM Torin2 reduced culture fresh weight by 50% over 14 days compared to untreated control, underscoring TOR’s role in sustaining cell growth and proliferation (Fig. 2a). On NLN medium ^34^ containing 1% (∼30 mM) sucrose, supplementation with 100 μM T6P—but not 100 μM sucrose—more than doubled fresh weight (Fig. 2B). This growth-promoting effect of T6P was largely abolished by the addition of 1 μM Torin2 (Fig. 2b). Metabolite analysis revealed that cells treated with T6P accumulated higher intracellular T6P at 3 and 8 h compared to sucrose-treated cells, while sucrose levels remained unchanged (Fig. 2c, d). Given that exogenous T6P (100 μM) is far below the sucrose concentration (30 mM), these findings indicate that T6P promotes cell proliferation primarily by activating TOR, supporting its role as a signaling metabolite distinct from sucrose metabolism.

**Fig. 2.**
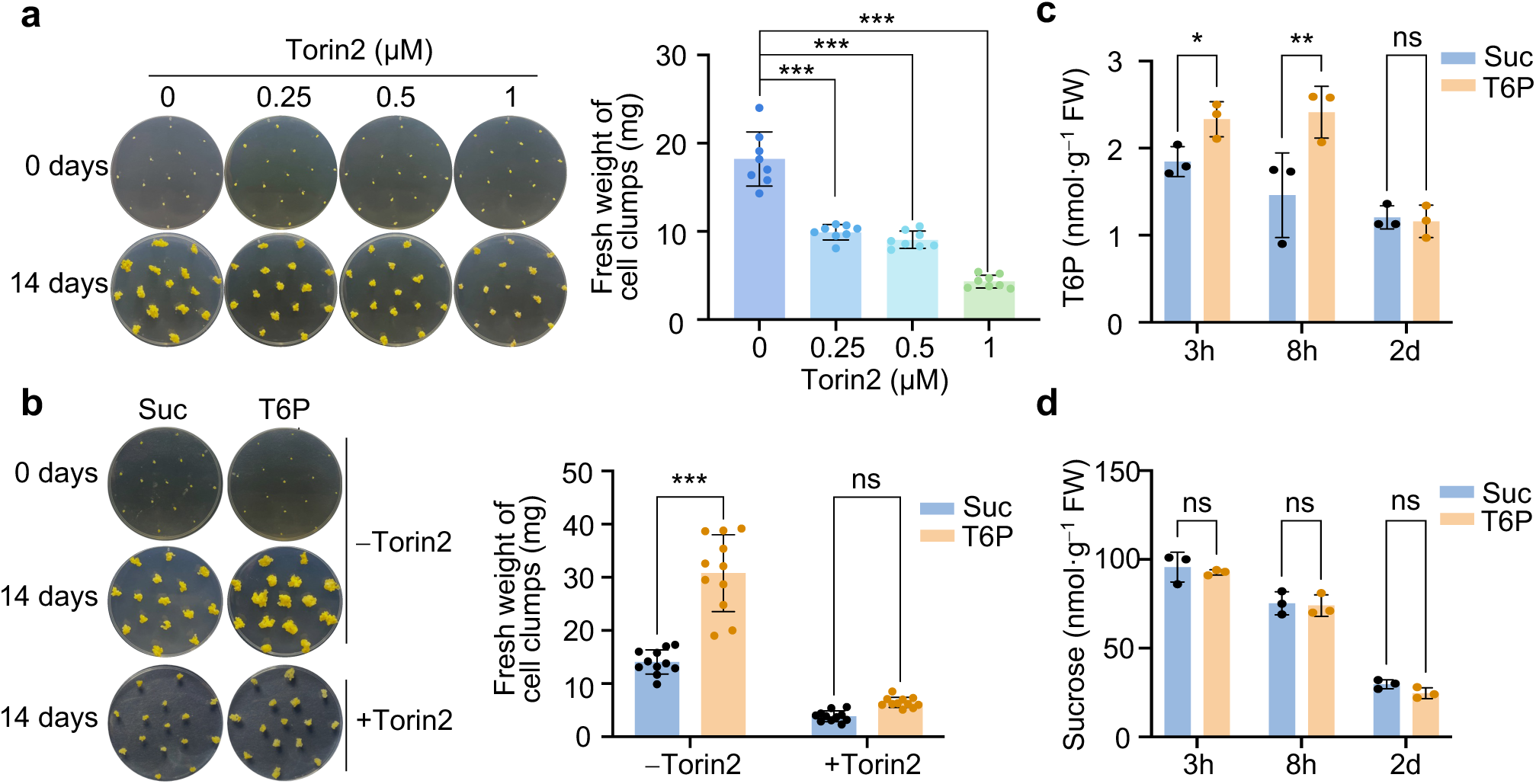
T6P promotes *Brassica napus* suspension cell growth via TOR. **a**, TOR activity is required for cell growth. Torin2 treatment markedly reduces biomass accumulation in *B. napus* suspension cells. Cell clumps (∼1 mm) were placed on NLN (1% sucrose) with 0–1 μM Torin2 and cultured at 28 °C for 14 d. Representative plates are shown. Plates are 4.5 cm in diameter. Fresh weight of cell clumps was measured after 14 d. **b**, T6P stimulates growth in a TOR-dependent manner. Exogenous T6P, but not sucrose, enhances biomass accumulation, and this effect is blocked by Torin2. Cells were grown on NLN (1% sucrose) supplemented with 100 μM sucrose or 100 μM T6P ± 1 μM Torin2 for 14 d. T6P (**c**) accumulates intracellularly after supplementation, while sucrose levels (**d**) remain unchanged. In all panels, significance by Tukey’s multiple comparisons: *, *P* < 0.05; **, *P* < 0.01; ***, *P* < 0.001, ns, not significant.

### T6P is essential for sucrose-mediated TOR activation *in vivo*

We next asked whether T6P functions as a signaling intermediate linking sucrose to TOR. TOR activity was monitored in Arabidopsis suspension cells derived from seedlings of an *S6K1-HA* overexpressing line^4^, using S6K1 phosphorylation and the pRPS6/RPS6 ratio as readouts. Exogenous T6P (100 μM) induced stronger TOR activation than equimolar sucrose during nutrient resupply (Fig. 3a), consistent with the *B. napus* growth results.

**Fig. 3.**
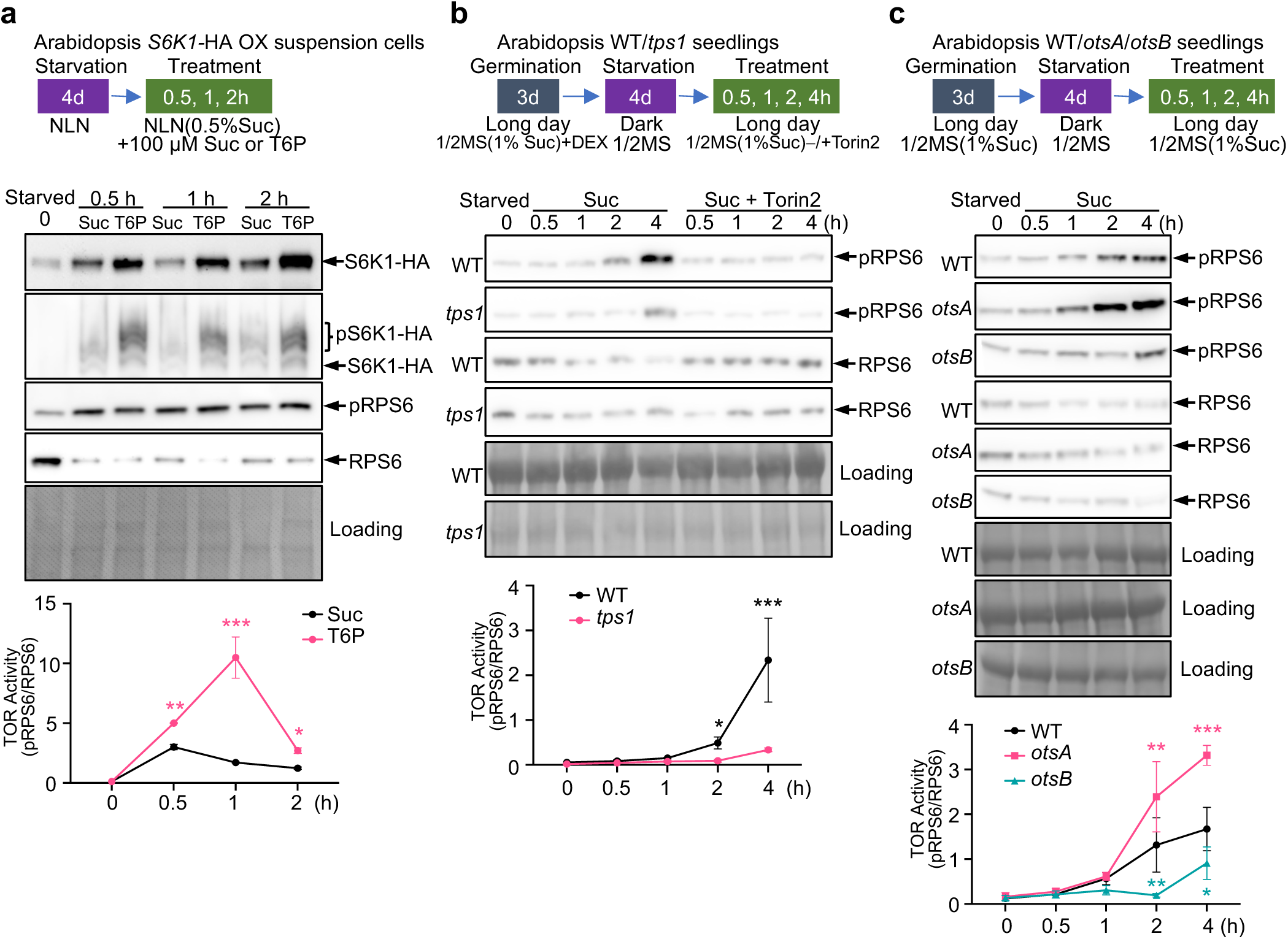
T6P is required for sucrose-mediated TOR activation. **a**, Exogenous T6P induces stronger TOR activation than sucrose during resupply of starved suspension cells. Starved suspension cells derived from an Arabidopsis *S6K1-HA* OX seedlings were resupplied with 0.5% of basal sucrose plus 100 μM sucrose or 100 μM T6P (0–2 h). Representative immunoblots show S6K1-HA, phosphorylated S6K1-HA (pS6K1-HA), phosphorylated RPS6 (pRPS6), and RPS6 at the indicated time points. Phos-tag gels separate phospho-S6K1-HA. Ponceau S serves as loading control. Band intensities were quantified using GelAnalyzer 23.1.1 and normalized to total protein. pRPS6/RPS6 ratios (n=3) were quantified for TOR activity (Šídák’s Multiple Comparisons Test: *, *P* < 0.05; **, *P* < 0.01; ***, *P* < 0.001). **b**, Endogenous T6P is essential for TOR activation. In WT seedlings, sucrose resupply robustly activates TOR, whereas *tps1* mutants show only weak activation. 3-d-old seedlings germinated on ½ MS (1% sucrose) (long-day conditions) were transferred to ½ MS liquid medium and starved for 4 d in dark. Seedlings were then resupplied with 1% sucrose ± 1 μM Torin2. Samples at 0.5–4 h. Representative immunoblots and quantified TOR activity (described as in **a**) are shown. **c**, Modulating T6P alters sucrose-mediated TOR activation. *otsA* accelerates and *otsB* delays TOR activation relative to WT. 3-d-old WT, *otsA* and *otsB* seedlings germinated on ½ MS (1% sucrose) (long-day conditions) were transferred to ½ MS liquid medium and starved for 4 d in dark. Seedlings were then resupplied with 1% sucrose. Samples at 0.5–4 h. Representative immunoblots and quantified TOR activity (described as in **a**) are shown.

To evaluate the requirement of endogenous T6P, we compared WT and *tps1* seedlings under sucrose resupply. In WT, TOR activity increased robustly at 2 hours (h) and continued to rise through 4 h. By contrast, *tps1* seedlings exhibited only modest TOR activation at 4 h (Fig. 3b). We also examined *otsA* and *otsB* overexpression lines. In starved seedlings, sucrose-induced TOR activation was significantly enhanced in *otsA* but delayed and attenuated in *otsB* relative to WT (Fig. 3c). These findings demonstrate that T6P is indispensable for sucrose-triggered TOR activation in vivo, establishing it as a critical mediator of the sugar–TOR signaling axis.

### T6P activates TOR kinase activity by suppressing SnRK1

Transcriptional studies showed that transient T6P elevation upregulates genes involved in protein and nucleotide biosynthesis, processes downstream of TOR^1, 35, 36^. However, the rapid TOR activation observed within minutes of T6P treatment in suspension cells (Fig. 2a) suggested a more rapid, possibly post-transcriptional regulatory mechanism.

To test this, we established an *in vitro* TOR kinase assay using TOR complexes immunoprecipitated from Arabidopsis seedlings (IP-TOR; Supplementary Fig. 6) with an antibody made against a C-terminal fragment of TOR (AA2282–2481)^35^. IP-TOR phosphorylated purified TOR substrates—human EIF4EBP1 or MBP-tagged Arabidopsis S6K1—within 15 min at 30 °C. Phosphorylation was absent when IP-TOR was omitted, and was abolished by Torin2 and enhanced by the addition of constitutively active ROP2 (CA-ROP2), a known TOR activator^8^, validating assay specificity (Fig. 4a, b, Supplementary Fig. 7).

**Fig. 4.**
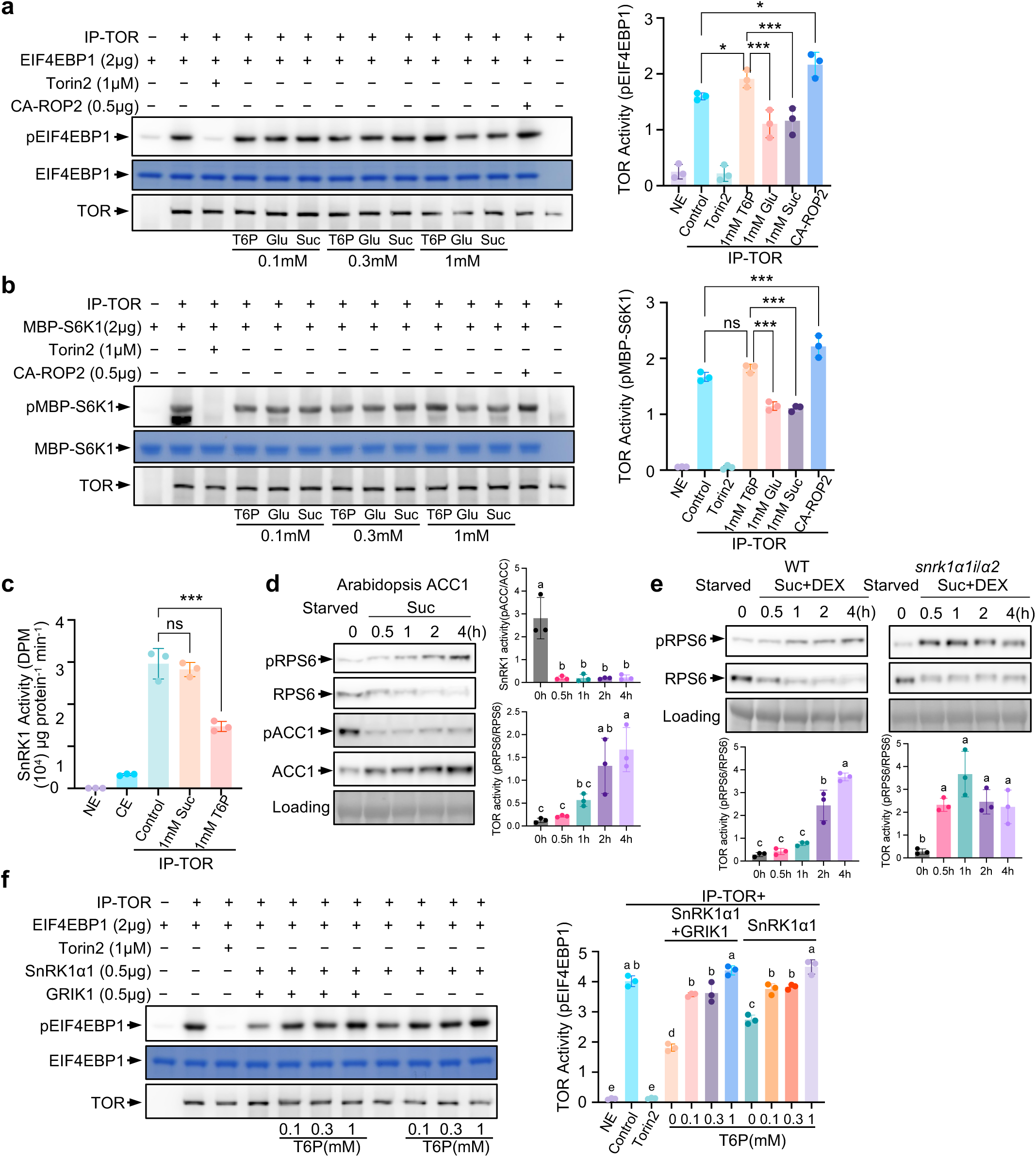
T6P activates TOR by suppressing SnRK1. *In vitro* TOR kinase assays show T6P (1 mM) modestly enhance TOR activity, where glucose or sucrose have little effect on TOR activity. Immunoprecipitated TOR (IP-TOR) from 7-d-old seedlings were incubated with EIF4EBP1 (**a**) or MBP-S6K1 (**b**) for 15 min at 30 °C. Reactions contained 0.1, 0.3 or 1 mM of T6P, glucose (Glu) or sucrose (Suc). Control, the basal reaction without supplementation. The reaction lacking IP-TOR (NE, no enzyme) or containing Torin2 served as negative controls; the reactions containing purified constitutively active ROP2 (CA-ROP2) served as a positive control. Phosphorylated EIF4EBP1 was detected with phospho-specific antibodies against EIF4EBP1 (pT37) and AtS6K1 (pT449), respectively. Equal substrate loading was verified by Coomassie, equal TOR by anti-TOR. Quantification at right. **c**, SnRK1 activity co-immunoprecipitates with TOR and is directly inhibited by T6P. AMARA-peptide assays with [γ-^32^P] ATP revealed SnRK1 activity in IP-TOR that was significantly reduced by 1 mM T6P versus 1 mM sucrose. NE, no enzyme. CE, crude extract. **d**, *In vivo*, sucrose resupply triggers reciprocal changes in TOR and SnRK1 activity. In a transgenic SnRK1-reporter line expressing a rat ACC1 peptide, SnRK1 activity rapidly declined as TOR activity gradually increased following sucrose resupply. Representative immunoblots show pRPS6, RPS6, pACC1, and ACC1. TOR and SnRK1 activities were quantified (right). Different letters above bars indicate statistically significant differences (Tukey’s multiple comparisons test, n=3, *P* < 0.05). **e**, *snrk1* inefficient mutant shows accelerated TOR activation during sucrose-mediated TOR activation. In the presence of DEX, in starved *snrk1α1i*/*α2* (DEX-inducible RNAi knockdown of *SnRK1α1* in a *SnRK1α2* knockout) seedlings, TOR demonstrated earlier activation after sucrose resupply compared to that in WT. TOR activities were quantified (below). **f**, Purified SnRK1α1 suppresses TOR activity, and T6P reverses this inhibition in a dose-dependent manner. IP-TOR incubated with SnRK1α1 alone or with GRIK1 and EIF4EBP1 substrate ± T6P (0.1–1 mM). TOR activities were quantified at right. Similar results were shown with S6K1 (Supplementary Fig. 8).

We then tested whether T6P directly stimulates TOR. Supplementing kinase reactions with 0.1 or 0.3 mM T6P, glucose or sucrose had no effect. At 1 mM, T6P significantly enhanced TOR activity compared to glucose or sucrose (Fig. 4a, b). However, the enhancement by 1 mM T6P was modest relative to no-sugar control—indistinguishable for S6K1 substrate and only marginal for EIF4EBP1 (P < 0.05) — suggesting that T6P does not strongly activate TOR directly.

Given that T6P is a known inhibitor of SnRK1^25, 26, 27, 28^ which has been reported to phosphorylate the RAPTOR subunit of the TOR complex^18^. We hypothesized that T6P potentiates TOR activity indirectly by suppressing SnRK1 associated with TOR complexes. Indeed, using AMARA peptide SnRK1 assay^25^, SnRK1 activity was detected in IP-TOR samples (Fig. 4c), consistent with a report that SnRK1α1 co-immunoprecipitates with TOR^22^. Addition of 1 mM T6P reduced SnRK1 activity in IP-TOR by >50% relative to the no-sugar or sucrose-supplemented controls (Fig. 4C), consistent with direct inhibition of SnRK1 by T6P. In a SnRK1-reporter line expressing a rat ACC1 peptide^37^, SnRK1 activity declined rapidly as TOR activity gradually increased following sucrose resupply, demonstrating reciprocal regulation (Fig. 4d). Consistently, TOR activation was markedly accelerated in the *snrk1α1i*/*α2* line (DEX-inducible RNAi knockdown of *SnRK1α1* in a *SnRK1α2* knockout background) ^37^compared to WT, as early as 0.5 h after sucrose resupply in the presence of DEX (Fig. 4e). To test whether T6P directly relieves SnRK1-mediated TOR inhibition, we supplemented the IP-TOR kinase assay with purified SnRK1α1 alone or activated with GRIK1, along with increasing concentrations of T6P. SnRK1α1 strongly inhibited TOR activity, consistent with the AMPK inhibition of mTOR in mammalian systems^38, 39^. Strikingly, T6P fully reversed this inhibition in a dose-dependent manner for both EIF4EBP1 and S6K1 substrates (Fig. 4f, Supplementary Fig. 8). Finally, microscale thermophoresis revealed that T6P binds IP-TOR with a dissociation constant (Kd) of 16.1 ± 2.46 μM (Supplementary Fig. 9), similar to its reported affinity for purified SnRK1α1^27^. By comparison, ATP bound TOR more tightly (Kd = 5.89 ± 0.83 μM).

Together, these findings demonstrate that T6P can activate TOR primarily by suppressing SnRK1, thereby releasing TOR from repression.

**Fig. 5.** A Model for T6P-mediated activation of TOR. Under favorable conditions, sucrose elevates T6P through TPS1. T6P negatively feeds back on sucrose and binds/inhibits SnRK1 (via SnRK1α1) preventing phosphorylation of RAPTOR and relieving TOR from repression. Activated TOR phosphorylates S6K1 and other effectors to stimulate biosynthesis, growth, and development.(Created with BioRender.com)

## Conclusion

T6P functions as a sucrose-proximal signal that activates TOR by suppressing SnRK1. This mechanism is particularly evident during sucrose-mediated TOR activation, positioning T6P as a key metabolite mediating the antagonistic TOR–SnRK1 relationship. By tightly coupling carbon status to growth decisions, this sucrose–T6P–SnRK1–TOR signaling axis provides a long-sought molecular link between metabolism and development, with broad implications for optimizing crop productivity and resilience in fluctuating environments.

## Materials and Methods

### Plant materials and growth conditions

*Arabidopsis thaliana* (Columbia-0) and the following lines were used: estradiol-inducible TOR RNAi (*tor-es1* and *tor-es2*; ABRC Stock# CS69829 and CS69830)^4^, hypermorphic TOR promoter insertion alleles (G548 and S7817; GK-548G07-020632 and salk_147817)^33^, S6K1-HA overexpression (ABRC Stock# CS73259) ^4^, *tps1-2; GVG::TPS1*^40, 41, 42^ (hereafter referred to as *tps1*), a dexamethasone (DEX)-inducible *TPS1* in a *tps1* knockout background, *snrk1α1i*/*α2* line, a DEX-inducible RNAi knockdown of *SnRK1α1* in a *SnRK1α2* knockout background^37^, and a SnRK1-reporter line expressing a rat ACC1 peptide (GFP-2×ACC1-2×HA)^37^.

For germination under long-day conditions, surface-sterilized seeds were sown on half-strength Murashige and Skoog (½ MS) medium with 1 % (w/v) sucrose and 0.4 % (w/v) phytagel. Seeds were stratified for 3 d at 4 °C in the dark and germinated in a Percival chamber (16 h light/ 8 h dark at 24 °C /22 °C, 150 μmol m⁻² s⁻¹).

For thermomorphogenesis assays, 3-d-old seedlings were transferred to fresh ½ MS (1% sucrose) and grown at 20 °C or 28 °C for 4 d under constant light (30 μmol m⁻² s⁻¹). For starvation/resupply assays, 3-d-old seedlings were transferred to ½ MS liquid medium, kept in darkness for 4 d, then resupplied with 1% sucrose with or without 1 μM Torin2, and sampled at 0.5, 1, 2 and 4 h.

### Brassica napus suspension cells and experimental treatments

Microspore-derived *B. napus* cv Jet Neuf suspension cells were propagated on Nitsch & Nitsch (NLN) solid media containing 3% sucrose and 0.8% agar for 7 d^43, 44^. Bright yellow cell clusters (∼ 1 mm) were transferred to NLN with 1% sucrose and 0.8% agar, supplemented with either 100 μM sucrose or 100 μM T6P. Torin2 was added where indicated. Plates were incubated at 28 °C under constant light (30 μmol m⁻² s⁻¹). Fresh weight was measured after 14 d.

### Metabolite measurements

Free metabolites were extracted as described in Mata et al. ^45^with modifications. Briefly, 50 mg of fresh plant material was homogenized in liquid nitrogen using a Geno/Grinder, extracted with 500 μL of ice-cold chloroform/methanol (3:7, v/v), vortexed, and incubated at −20 °C for 2 h with intermittent vortexing. After adding 400 μL ice-cold water, samples were incubated at 4 °C for 15 min. Deoxy-G6P was included as internal standard. Extracts were centrifuged (14,000 rpm, 10 min, 4 °C), and then aqueous phase was collected, re-extracted, pooled, and freeze-dried. T6P was quantified using a two-step derivatization method^46^, Freeze-dried samples were dissolved in 100 µL of methoxylamine (20 mg/mL in pyridine) and incubated at 60 °C for 30 min, then overnight at room temperature. Next, 30 µL of 1-methylimidazole and 60 µL of propionic acid anhydride were added and incubated at 37 °C for 30 min. The mixture was dried with nitrogen and re-dissolved in 100 µL of 0.1% formic acid. LC-MS used a Shimadzu Nexera X2 UHPLC system coupled to a Sciex QTRAP 4500. Derivatized sugar phosphates were separated on an EVO C18 column (1.7 μm, 100 Å, 100 × 2.1 mm, Phenomenex, Torrance, CA) in negative ionization mode.

Sucrose was analyzed on a Luna NH₂ column (100 × 2 mm, 3 μm; Phenomenex) in negative ion mode. Solvents: A, acetonitrile; B, 20 mM ammonium acetate/20 mM ammonium hydroxide in 95:5 water: acetonitrile (pH 9.45). Gradient: 25% B (0 min) → 30% B (8 min) → 100% B (22–32 min) → 25% B (33.5–44 min). Flow, 0.3 mL min⁻¹; column, 40 °C. Quantification used ¹³C₆-glucose as internal standard.

### Production and purification of recombinant protein from E. coli

Coding sequences of KIN10, GRIK1, S6K1, EIF4EBP1, CA-ROP2 (Q63L) were cloned into the pET28a or pDEST-HisMBP vector using In-fusion^TM^ cloning. N-terminal His-tag proteins were expressed in *E. coli* BL21 (DE3) and purified as described^47^.

### Immunoblotting

Immunoblotting followed established protocols^48^. Seedlings (20 mg) were ground in liquid nitrogen and extracted in 80 µL buffer [8 M urea, 2 % (w/v) SDS, 0.1 M DTT, 20% (v/v) glycerol, 0.1 M Tris-HCl, pH 6.8, and 0.004 % Bromophenol Blue]. Extracts were heated (80 °C, 5 min), centrifuged (17,000 ×g), and 10 μL of supernatant was resolved by SDS-PAGE. Proteins were transferred to a PVDF membrane, blocked with 5 % (w/v) milk, and probed with primary antibodies: anti-TOR (1:3,000), anti-HA (1:3,000; Catalog# 901514, Biolegend), anti-phospho-RPS6 (1:5,000; a gift from Prof Christian Meyer, IJPB)^5^, anti-RPS6 (1:1,000; Catalog# 2317, Cell Signaling Technology). HRP-conjugated secondary antibodies (1:10,000; Catalog# AS09 602, Agrisera) were detected using SuperSignal^TM^ West Femto Maximum Sensitivity Substrate (Catalog# 34095, ThermoFisher) on an Amersham ImageQuant 800 (Cytiva).

Phosphorylated proteins were resolved by Phos-tag SDS-PAGE (SuperSep^TM^ Phos-tag^TM^ (50 μmol/L), 10 %, Catalog# 193-16711, FUJI Film). After electrophoresis, gels were washed three times in the protein transfer buffer (25 mM Tris, 0.192 M Glycine, 10 % (v/v) methanol) containing 10 mM EDTA, before transfer to a PVDF membrane.

### Immunoprecipitation (IP) of native TOR

Native TOR was immunoprecipitated as described^49^. 1 gram of 7-d-old WT seedlings grown on ½ MS (1% sucrose) was ground in liquid nitrogen and extracted in 5 mL of protein extraction buffer (400 mM HEPES pH 7.5, 2 mM EDTA, 10 mM Pyrophosphate, 10 mM Glycerol phosphate, 0.3 % (w/v) CHAPS, protease inhibitor cocktail (cOmplete^TM^, Mini EDTA-free, Catalog# 04693132001, Roche). Extracts were rotated 15 min at 4 °C, clarified by sequential centrifugation (20,000 ×g) and filtered through Econo-Pac® Disposable Chromatography Column. The supernatant was concentrated using a 100 kDa Amicon Ultra Filter (Merck Millipore, Billerica, MA, USA) via centrifugation at 3,500 ×g. Dynabeads® Protein A (Catalog# 10001D, ThermoFisher Scientific) were conjugated with anti-TOR antibody and incubated with protein extracts (2 h, 4 °C). Complexes were captured magnetically, washed once in protein extraction buffer and twice in low-salt buffer(400 mM HEPES pH 7.5, 150 mM NaCl, 2 mM EDTA, 10 mM Pyrophosphate, 10 mM Glycerol phosphate, 0.3 % (w/v) CHAPS), then resuspended in kinase wash buffer (25 mM HEPES pH 7.5, 20 mM KCl).

### TOR in vitro kinase assay

Reactions (25 μL) contained 25 mM HEPES (pH 7.5), 50 mM KCl, 10 mM MgCl_2_, 0.02 mM ATP, 5 μL of immunoprecipitated TOR (on beads), and 2 μg EIF4EBP1 or MBP-AtS6K1 substrate protein. Reactions were incubated at 30 °C for 15 min and terminated with SDS-PAGE buffer. TOR activity was assessed by phosphorylation detected with phospho-specific antibodies against EIF4EBP1 (pT37/46) (Catalog# 2855, Cell Signaling Technology) or AtS6K1 (pT449) (Catalog#AS132664, Agrisera). To assess sugars effects, TOR-bead complexes were preincubated (10min, on ice) with T6P, glucose or sucrose (100 μM–1 mM). Negative controls lacked TOR or included 1 μM Torin2; positive controls included purified constitutively active ROP2 (CA-ROP2).

### SnRK1 kinase assay

Crude protein extracts were prepared in buffer as above. Ten µg crude extract protein or 2 µg IP-TOR was incubated with 200 μM AMARA peptide in kinase assay buffer (25 mM HEPES pH 7.5, 50 mM KCl, 10 mM MgCl₂, 20 μM cold ATP, 2 µCi [γ-^32^P] ATP (Catalog# BLU002Z250UC, Revvity) at 30 °C for 10 min. Reactions were spotted (5 µl) on 4 cm^2^ phosphocellulose paper (ion exchange cellulose chromatography paper, Lab Alley), washed five times in 1% (v/v) phosphoric acid, rinsed in acetone, dried, and scintillation-counted (Hidex)^27,50^. Activity is expressed as pmol phosphate incorporated per min per mg protein. Each sample was assayed in triplicate.

### Microscale Thermophoresis (MST)

MST was performed using a Monolith NT.115 (NanoTemper Technologies, South San Francisco, CA). IP-TOR was fluorescently labeled with NT647 dye (NanoTemper Technologies) via N-hydroxysuccinimide coupling; unbound dye was removed by size-exclusion chromatography in MST buffer [50 mM phosphate buffer, pH 7.4, 150 mM NaCl, 10 mM MgCl_2_, 1 mM DTT, and 0.05 % (v/v) Tween 20]. For the binding assays, 200 nM labeled protein was incubated with 25 μM T6P (Catalog# 136632-28-5, Sigma) for 10 min before measurement. To determine the dissociation constants (*K_d_*), labeled protein was incubated with serial dilutions of T6P. 10 μL per sample was loaded into MST capillaries. Measurements were at 25 °C (40 % MST power and 80 % LED power).

## Accession Numbers

AT1G50030 (TOR), AT3G01090 (KIN10), AT1G20090 (ROP2), AT3G45240 (GRIK1), AT3G08730 (S6K1), Q13541 (EIF4EBP1).

## Acknowledgement

We thank Markus Schmid and Eunkyoo Oh for *tps1-2; GVG::TPS1*, Takeo Sato for SnRK1-reporter line (ACC1) and *snrk1α1i*/*α2*, and Christian Meyer for antibodies against phosphorylated RPS6 and RPS6.

## Author Contributions

Conceptualization: Z. Z., J. Sh.

Methodology: H. L., J. B., H. S., S. A., J. Sc., J. Sh., Z. Z.

Investigation: H. L., J. B., H. S., S. S., S. A., J.Sc., J. Sh., Z. Z.

Visualization: H. L., J. B., J. Sh., Z. Z.

Funding acquisition: Z. Z., J. Sh.

Project administration: Z. Z., J. Sh.

Supervision: Z. Z., J. Sh.

Writing – original draft: Z. Z.

Writing – review & editing: Z. Z., J. Sh.

## Data and materials availability

All data are available in the Manuscript or the Supplementary Materials. Z. Z. is responsible for the distribution of materials integral to the findings.

