## Supplementary Figures for "Trehalose 6-phosphate activates Target of Rapamycin in plants"

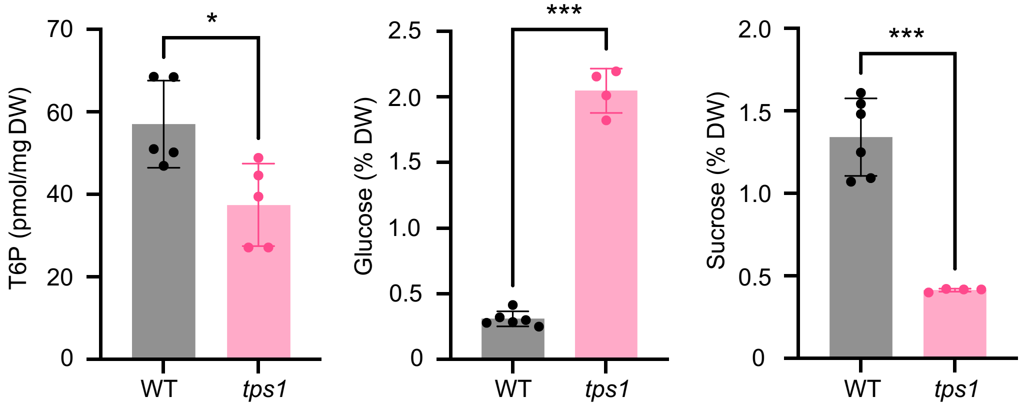


**Supplementary Fig. 1. T6P content is reduced in *tps1* seedlings without dexamethasone.**
T6P, glucose, and sucrose levels were quantified in WT and *tps1* seedlings grown on ½ MS with 1% sucrose. *tps1* seedlings show significantly reduced T6P, confirming that the system effectively lowers endogenous T6P. Student's t-test: *, $P < 0.05$; **, $P < 0.01$; ***, $P < 0.001$.

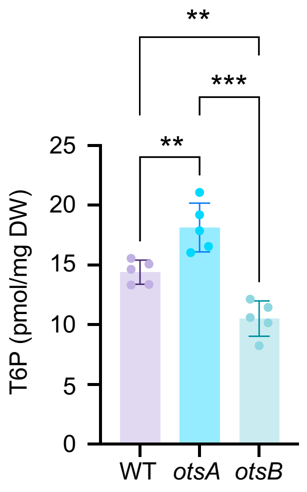


**Supplementary Fig. 2. T6P levels in WT, *otsA*, and *otsB* lines.**
Leaf tissues from 20-day-old plants were analyzed for T6P content. *otsA* overexpression increases T6P, whereas *otsB* overexpression reduces T6P, validating these lines for functional studies. Student's t-test: *, $P < 0.05$; **, $P < 0.01$; ***, $P < 0.001$.

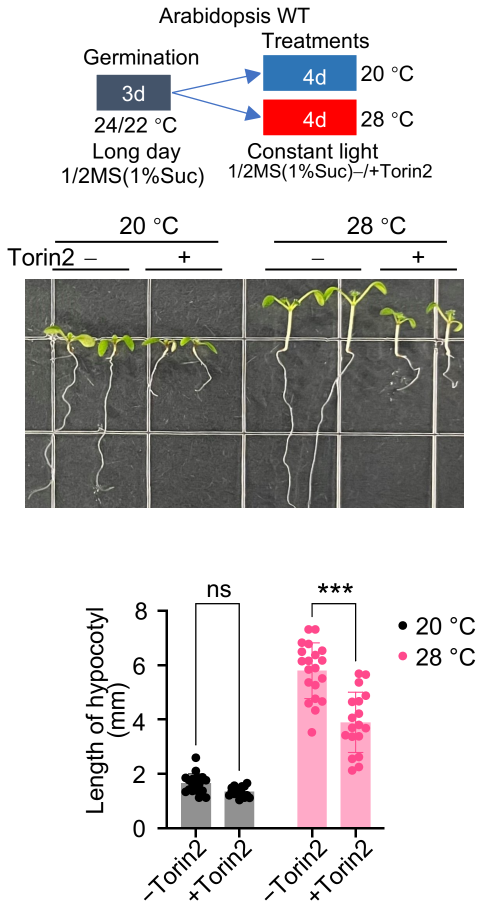


**Supplementary Fig. 3. TOR inhibition reduces plant thermoresponses.** WT seedlings grown at 28 °C in the presence of Torin2 show shorter hypocotyls compared with untreated controls. This confirms that TOR activity is necessary for thermomorphogenesis. WT seeds were germinated on ½ MS (1% sucrose) for 3 d (long-day conditions), then transferred to ½ MS (1% sucrose) ± 1 μM Torin2 and grown at 20 °C or 28 °C for 4 d under constant light. Representative seedlings and quantified hypocotyl lengths are shown. Grid size, 13.5 × 13.5 mm. Significance by Tukey's multiple comparisons: *, $P < 0.05$; **, $P < 0.01$; ***, $P < 0.001$, ns, not significant.

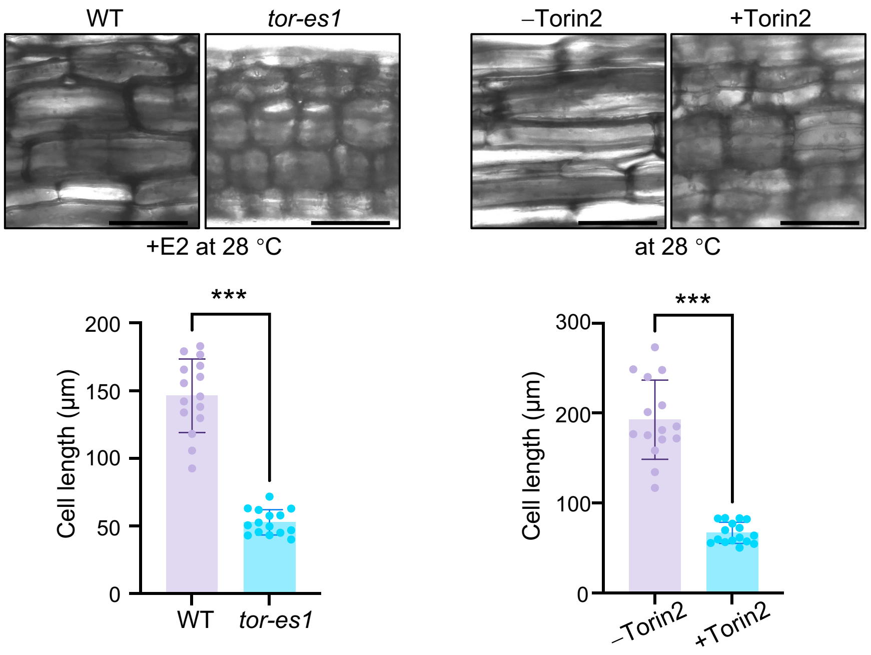


**Supplementary Fig. 4. TOR inhibition reduces epidermal cell length in Arabidopsis hypocotyls.**

Differential interference contrast imaging reveals shorter epidermal cells in TOR-silenced (*tor-es1*) or Torin2-treated seedlings relative to controls. WT or *tor-es1* seedlings grown at 28 °C with estradiol (E2), or from WT seedlings grown at 28 °C without or with 1 µM Torin2 for 4 d. Scale bar, 100 µm. Corresponding epidermal cell lengths are shown. Student's t-test: *, $P < 0.05$; **, $P < 0.01$; ***, $P < 0.001$.

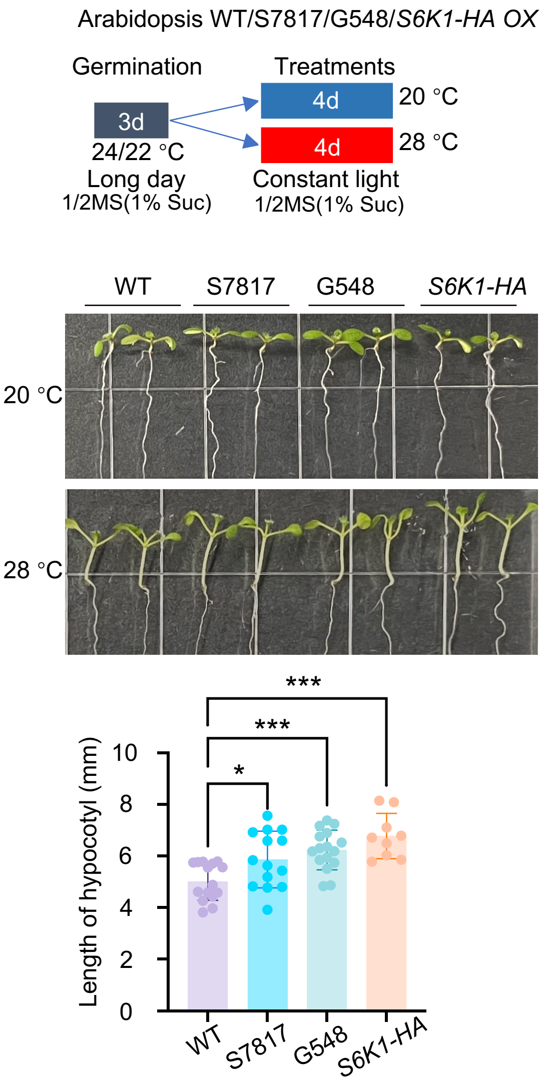


**Supplementary Fig. 5. TOR or S6K1 overexpression enhances thermoresponsive growth.**

Seedlings carrying hypermorphic TOR alleles or overexpressing S6K1 display longer hypocotyls at 28 °C compared with WT. Elevated TOR signaling therefore enhances thermomorphogenesis. WT, TOR hypermorphic alleles (S7817, G548) and S6K1-HA overexpression lines were grown as indicated; representative seedlings and quantified hypocotyl lengths (28 °C) are shown. Student's t-test: *, $P < 0.05$; **, $P < 0.01$; ***, $P < 0.001$.

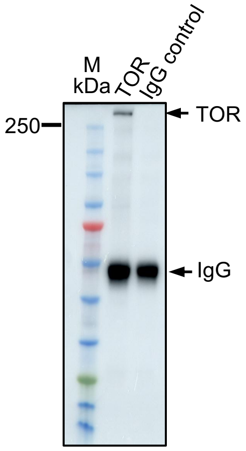


**Supplementary Fig. 6. Immunoprecipitated TOR from Arabidopsis seedlings.**
TOR was successfully immunoprecipitated using a custom antibody; pre-immune IgG did not recover TOR. This validates the specificity of the IP-TOR complex used in kinase assays.

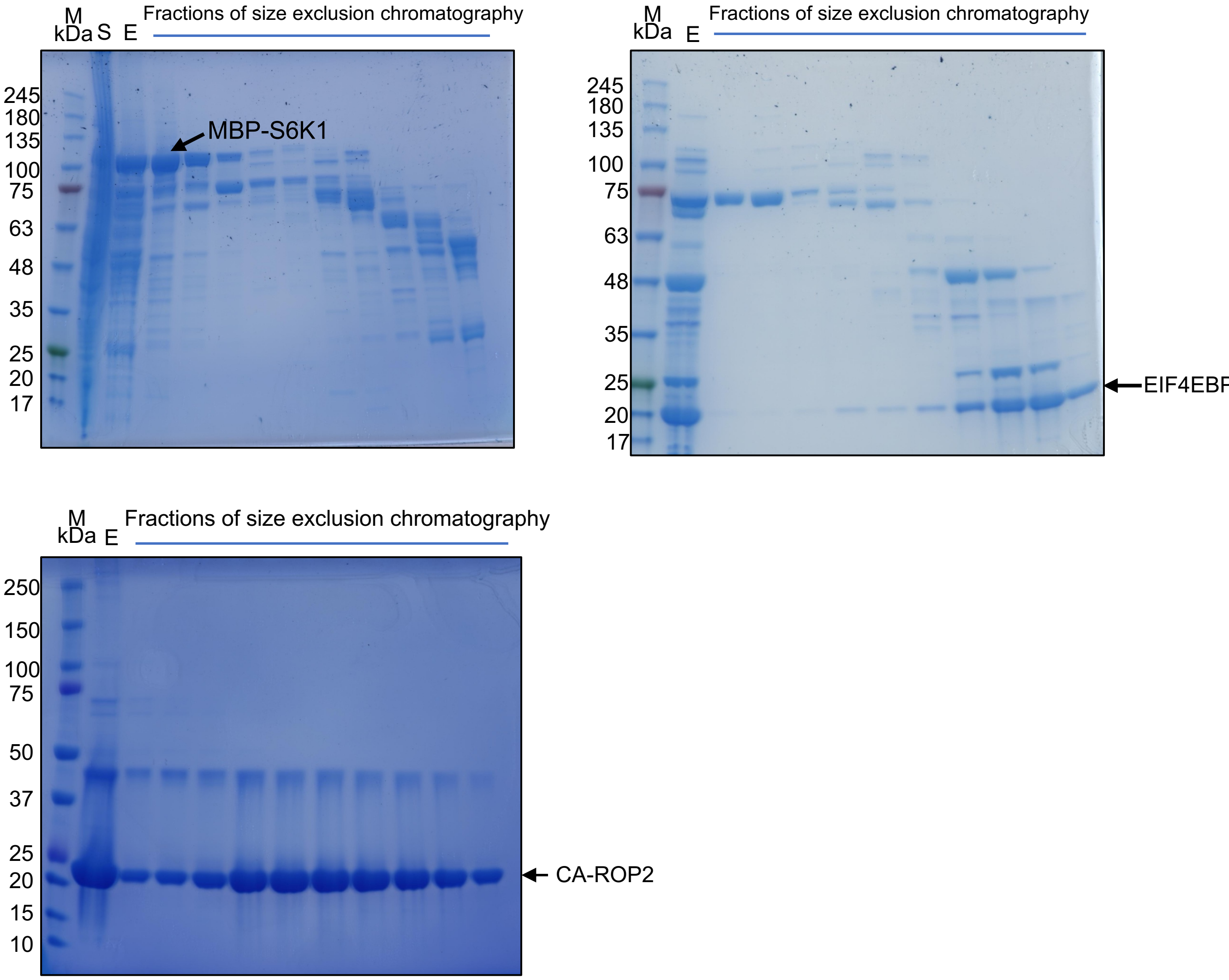


**Supplementary Fig. 7. Purification of recombinant proteins for TOR kinase assays.** Coomassie-stained gels confirm the successful expression and purification of MBP–S6K1, EIF4EBP1, and CA-ROP2 proteins from *E. coli*. M, molecular weight marker; S, supernatant after centrifugation of the cell lysate; E, elution from Ni–NTA affinity column.

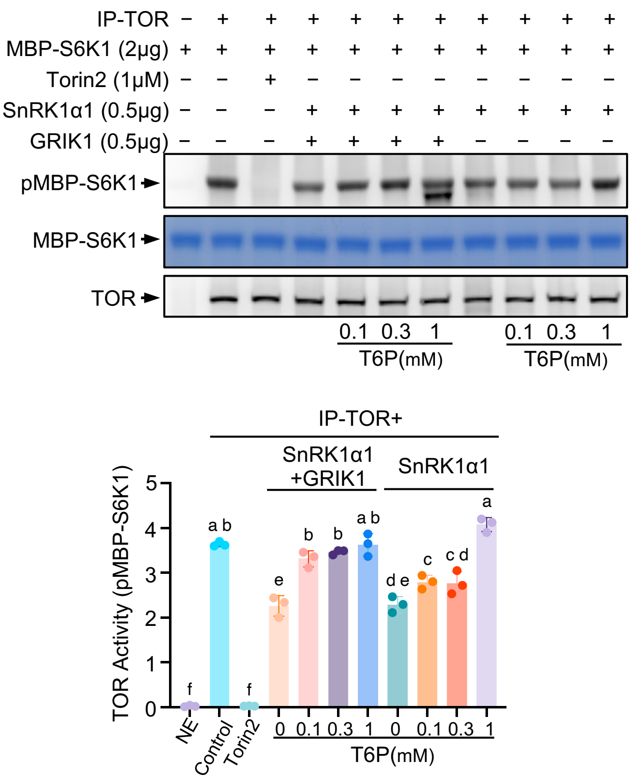


**Supplementary Fig. 8. T6P relieves SnRK1-mediated inhibition of TOR *in vitro*.** TOR kinase assays show that SnRK1α1 strongly inhibits TOR activity, and increasing concentrations of T6P progressively restore TOR activity. This demonstrates that T6P can antagonize SnRK1 repression of TOR. IP-TOR was incubated with SnRK1α1 alone or with GRIK1 and S6K1 substrate ± T6P (0.1–1 mM). Different letters above bars indicate statistically significant differences (Tukey's multiple comparisons test, $P < 0.05$).

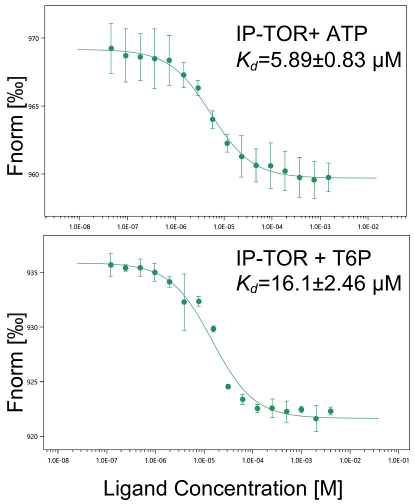


**Supplementary Fig. 9. ATP and T6P bind IP-TOR.**
Microscale thermophoresis (MST) measurements show that T6P binds immunoprecipitated TOR with micromolar affinity, comparable to prior reports for purified SnRK1α1 while ATP binds more tightly. These results confirm direct interaction of T6P with TOR complexes. Data are mean ± SD, $n$ = 3 biological replicates. Fnorm, normalized fluorescence (‰).
